# Comparison of morphine, oxycodone and the biased MOR agonist SR-17018 for tolerance and efficacy in mouse models of pain

**DOI:** 10.1101/2020.10.16.341776

**Authors:** Fani Pantouli, Travis W. Grim, Cullen L. Schmid, Agnes Acevedo-Canabal, Nicole M. Kennedy, Thomas D. Bannister, Laura M. Bohn

## Abstract

The mu opioid receptor-selective agonist, SR-17018, preferentially activates GTPγS binding over βarrestin2 recruitment in cellular assays. In mice, SR-17018 stimulates GTPγS binding in brainstem and produces antinociception with potencies similar to morphine. However, it produces much less respiratory suppression and mice do not develop antinociceptive tolerance in the hot plate assay upon repeated dosing. Herein we evaluate the effects of acute and repeated dosing of SR-17018, oxycodone and morphine in additional models of pain-related behaviors. In the mouse warm water tail immersion assay, an assessment of spinal reflex to thermal nociception, repeated administration of SR-17018 produces tolerance as does morphine and oxycodone. SR-17018 retains efficacy in a formalin-induced inflammatory pain model upon repeated dosing, while oxycodone does not. In a chemotherapeutic-induced neuropathy pain model SR-17018 is more potent and efficacious than morphine or oxycodone, moreover, this efficacy is retained upon repeated dosing of SR-17018. These findings demonstrate that, with the exception of the tail flick test, SR-17018 retains efficacy upon chronic treatment across several pain models.

## Introduction

Morphine-induced antinociception is mediated via the mu opioid receptors (MOR) that are expressed in key areas for pain control, in both spinal and supraspinal neurons (Bodnar, 2000; Kline and Wiley, 2008; Manning et al., 1994; Moriwaki et al., 1996; Narita et al., 1999; Yaksh, 1997). Cumulative evidence spanning three decades supports an involvement of the MOR in regulating pain responses to thermal (Yaksh, 1997), chemical (Manning et al., 1994) or mechanical nociceptive stimuli (Due et al., 2012; Johnson et al., 2014). As a G protein-coupled receptor (GPCR), the MOR signals through G proteins and is regulated by βarrestins.

βarrestin2 negatively regulates GPCRs including MOR; it was therefore proposed that removing this negative regulator of MOR signaling could enhance behavioral responses to a MOR agonist. Early studies using genetically modified mice lacking βarrestin2 demonstrated that morphine-induced antinociception was enhanced and prolonged in thermal assays of nociception (hot plate and warm water tail immersion assays) (Bohn et al., 2000; Bohn et al., 2002; Bohn et al., 1999). Further, the knockdown or deletion of βarrestin2 has been shown to attenuate morphine tolerance in rodent models (Bu et al., 2015; Li et al., 2009; Yang et al., 2011). We found that while βarrestin2-knockout (KO) mice did not become tolerant in the hot plate test, they did become tolerant in the warm water tail immersion assay (Bohn et al., 2002). A systemic injection of a nonselective protein kinase C (PKC) inhibitor, chelerythrine, was effective in reversing morphine tail flick tolerance in the βarrestin2-KO mice, but not in the WT mice (Bohn et al., 2002); these observations suggest that in the absence of βarrestin2, the contribution of other regulatory mechanisms, such as those involving PKC, becomes more apparent. These interpretations are limited by the fact that βarrestin2 is a ubiquitously expressed regulatory protein and therefore, its genetic deletion is likely to affect many regulatory and signaling cascades underlying nociceptive responses and drug adaptations upon chronic exposure.

Recently, we described a series of MOR agonists that preferentially induce MOR-mediated GTPγS binding over βarrestin2 recruitment (Schmid et al., 2017). Of these, SR-17018, was shown to be highly biased for G protein signaling (GTPγS binding assays in mMOR- and hMOR-expressing CHO cells, GTPγS binding mouse brainstem, cAMP accumulation studies in mMOR, hMOR CHO cells relative to βarrestin2 recruitment determined by enzyme fragment complementation to hMOR and βarrestin2-GFP translocation to mMOR), and was efficacious in antinociception while producing very little respiratory suppression. We demonstrated that SR-17018 had a similar potency as morphine in mouse brainstem and periaqueductal grey for stimulating GTPγS binding as well as in antinociception; moreover both effects were absent in MOR-KO mice (Schmid et al., 2017) (Grim et al., 2019). These observations together with demonstration that the compound is selective for MOR over other opioid receptors, suggests that SR-17018 is indeed acting at MOR *in vivo*. SR-17018 is orally bioavailable, has a 3:1 brain to plasma ratio of distribution and a pharmacokinetic half-life of 6-8 hours; moreover, it can be orally delivered to maintain consistent plasma levels over 6 days (Grim et al., 2019). Recently we showed that chronic oral administration (b.i.d., 6 days) with SR-17018 does not produce tolerance in the hot plate antinociception assay (Grim et al., 2019), which is in agreement with the early observations in the βarrestin2-KO mice (Bohn et al., 2002; Raehal and Bohn, 2011). We previously reported that treatment with SR-17018 could reverse morphine tolerance in the hot plate test while preventing the onset of withdrawal (Grim et al., 2019). Therefore, substitution with an agonist like SR-17018 may represent a means to restore MOR responsiveness and analgesic efficacy in the opioid dependent state, while lessening the severity of opiate withdrawal (Grim et al., 2019). In this study, we have further evaluated the efficacy of SR-17018 in acute and chronic treatment paradigms across different mouse pain models.

## Methods

### Animal Care and Use

Approximately 80% of the C57BL/6J mice used were acquired from The Jackson Laboratories, and the remaining 20% were generated by breeding at Scripps Florida. A total of 316 male C57BL/6J mice and 12 female mice (supplemental Figure 1) were used to complete these studies. A total of 107 male DBA/2J mice have been used for the completion of the paclitaxel-induced neuropathic pain study, as this strain has been reported to exhibit a robust responses in the neuropathic pain model and we hoped to reduce the number of animals needed by improving the response window (Smith et al., 2004); DBA mice were acquired from The Jackson Laboratories. All mice used were 10-20 weeks of age at the time of testing. For chronic treatment studies, mice were single housed prior to minipump implantation (to avoid pump contact between implanted mice) or twice daily oral gavage (to match morphine). Mice were kept on 12-hour light-dark cycle and had *ad libitum* access to standard rodent chow and water throughout testing procedures. The tail flick response data was obtained from assessing responses prior to performing hot plate responses in an effort to reduce the overall number of mice needed for the study; hot plate responses have been published (Grim et al., 2019). Male and female mice were used separately, and the data is presented accordingly. All studies were performed in mice that had not been used in prior studies. All mice were used in accordance with the National Institutes of Health Guidelines for the Care and Use of Laboratory Animals with the approval by The Scripps Research Institute Animal Care and Use Committee.

### Blinding

Investigators were blinded to identification of compound components in all studies. Details on the compound treatment blinding approach for the studies involving the tail flick, the use of osmotic minipumps (o.m.p) and per os (PO) dosing can be found as described for the hot plate assessments previously published (Grim et al., 2019). In brief, compound doses and vehicles were prepared as 10X stocks and labeled by key (A, B, C, etc.) on the day of the study by an independent lab member and mice were treated by key designation. All treatment groups were randomized independent of baseline responses. Responses were assessed by investigators blinded to acute and chronic treatments.

### Chemicals and compounds

Morphine sulfate pentahydrate, oxycodone HCl and buprenorphine HCl, were dosed relative to salt weight and were purchased from Sigma Aldrich or received from the NIDA Drug Supply. SR-17018 mesylate salt was synthesized as previously described (Schmid et al., 2017), and dosed relative to the base weight. Vehicle in this study refers to a 1:1:8 vehicle (10%DMSO, 10%Tween-80, 80%purified water) and it is used to dissolve all compounds unless specified. SR-17018 was validated by NMR and purity was greater than 95%. Dimethyl sulfoxide (DMSO), Tween-80, and 0.9% saline were purchased from Fisher. DMSO and Tween-80 were stored at 4°C in small volumes to prevent hydration and oxidation upon repeated opening of bottles. Chelerythrine (Fisher) is prepared in 5% DMSO in sterile water. Formaldehyde, 37% microfiltered was purchased from Electron Microscopy Science (VWR) and used as 100% formalin in a ratio of 1:20 (formalin: purified water). Paclitaxel (Tocris) is dosed relative to salt weight of the drug, prepared in 10% Cremophor ^®^ EL (Sigma Aldrich) in 0.9% saline and sonicated prior to administration.

### Warm water tail immersion response latency (tail flick assay)

Mice were gently held in a towel and the tip of tail (~2cm) was dipped into warm water maintained at 49°C by a circulating water bath heater and the latency to withdraw the tail was recorded with an upper limit of 30 seconds (Schmid et al., 2017). A maximum possible effect was calculated for each animal as %MPE= 100% x (after drug latency - baseline / 30 second limit-baseline). The average response time for the male C57BL/6J mice used in this assay was: 2.8 (2.6-2.9) seconds (n= 110 mice; mean with 95% CI).

### Hot plate response latency

Mice were tested as previously described (Grim et al., 2019). The hot plate is held at 52°C and a cut off time of 20 seconds is imposed. Responses are measured when mouse withdraws any paw from the surface, aside from walking. The average response time for the male C57BL/6J mice used in the chelerythrine reversal study was: 5.5 (4.8-6.2) seconds (n= 10 mice; mean with 95% CI).

### Formalin-induced paw licking

For the dose response studies, mice were pretreated with vehicle or compound for 45-minutes, during which time they were habituating to square acrylic enclosures that are sitting on mirrors for at least 30-minutes. Then mice were injected with 25 μL of 5% formalin in the plantar surface of the right hind paw using a 30 ½ gauge needle attached to a Hamilton syringe. Mice were then placed in new square acrylic enclosures and the time spent licking or biting the injected paw was recorded in 5 minute bins for 40 minutes by observers blinded to treatment conditions at the time of testing (Tarselli et al., 2011). The data were summed in bins separated into Phase 1 (0-10 minutes) and Phase 2 (16-40 minutes) to reflect the direct activation of nociceptors and subsequent inflammatory and/or central sensitization effects, respectively (Dubuisson and Dennis, 1977; Mogil et al., 1996; Tjolsen et al., 1992). To determine ED_50_, data were normalized to vehicle response by setting the bottom as 0 seconds and the top as the average seconds for vehicle treatment within the phase. The % max inhibition was determined by Y=100-(Y*100) where y=normalized data; the normalized data were fit to hyperbolic nonlinear regression where the max was constrained to 100% to derive the ED_50_.

### Paclitaxel-induced neuropathic pain model and von Frey measures of allodynia

Baseline mechanical sensitivity was assessed with the IIDC Life Science Electronic von Frey Anesthesiometer on two consecutive days prior to drug treatments as previously described (Martinov et al., 2013). Mice were placed in plexiglass chambers elevated on a standard large stand (base 36” X 16”) with black anodized mesh (31 /14” X 11”) from IITC Life Science with 1/4” size waffle holes to afford access to the plantar surface of the hind paws. Three individual, acrylic animal enclosures are placed on top of mesh allowing unobstructed view of the mice during testing. The rigid tip at the handle of the anesthesiometer was used to apply pressure to the plantar surface of the hind paw with gradually increasing intensity until paw retraction was observed and the maximum force reading (in grams, g) was recorded. Following 30-minutes acclimation in the chambers, baseline measurements were taken five times on alternating paws for each mouse and then averaged to determine baseline.

Baselines were collected on two consecutive days and mean of baseline per individual mouse was averaged for the two days. Mice were injected with paclitaxel on four alternating days with a dose of 1 mg/kg/day IP as previously described (Smith et al., 2004). Von Frey response measures were again taken on post-injection Day 7 (following paclitaxel administration), Day 11 and Day 14 in the morning to test for mechanical sensitivity. Mice were tested for allodynia 1 hour after drug treatment in the afternoon of Day 14. Two testing criteria were applied per individual mouse: the average initial baseline response latency must be above 2g; the ΔForce threshold must be greater than 1 g when comparing Day14 response to baseline (indicates paclitaxel induced neuropathic pain). In the dose response study, 1 mouse was eliminated based upon these criteria. Mice that passed the exclusion criteria were evenly distributed into drug treatment groups by an investigator not involved with measuring responses.

### Drug dosing for tolerance

*SR-17018* was administered orally (PO) for chronic treatment as repeated IP or subcutaneous dosing at high concentrations leads to precipitation at the site of injection (Grim et al., 2019). Gavage was administered using flexible plastic feeding tubes mounted on the end of a 1 mL syringe at a volume of 10 μL/g of body weight. SR-17018 is orally bioavailable (Grim et al., 2019) and is potent in the tail flick response: ED_50_: 11.1 *(6.4-19.0)* mg/kg, n=5 at 6, 12; n=8 at 24 mg/kg, PO, (SFig 1A). *Oxycodone* was used to match SR-17018 oral gavage dosing due to oral bioavailability and daily drug preparation was done as previously described (Grim et al., 2019). SR-17018 and oxycodone were administered in vehicle with half the dose PO every 12 hours for 6 full days. *Morphine* was dissolved in saline or saline was administered by osmotic minipump. Primed minipumps (200 μl reservoir, 1 μl/hour flow rate, ALZET model 2001), were filled according to individual mouse body weight (23-32 grams at the time of surgery) in saline as previously described in detail (Grim et al., 2019). Challenge doses of compound were administered 12 hours after the last PO, b.i.d. treatment.

To determine tolerance and cross tolerance, all acute treatments and challenges (following chronic treatment) were identical to the hot plate assessments as previously described (the tail flick responses from the animals in Figure 1 are presented along with the hot plate data from the prior report) (Grim et al., 2019). For the formalin studies, only single injections (IP) were made following chronic treatment paradigms and responses were recorded over time.

**Figure 1.**
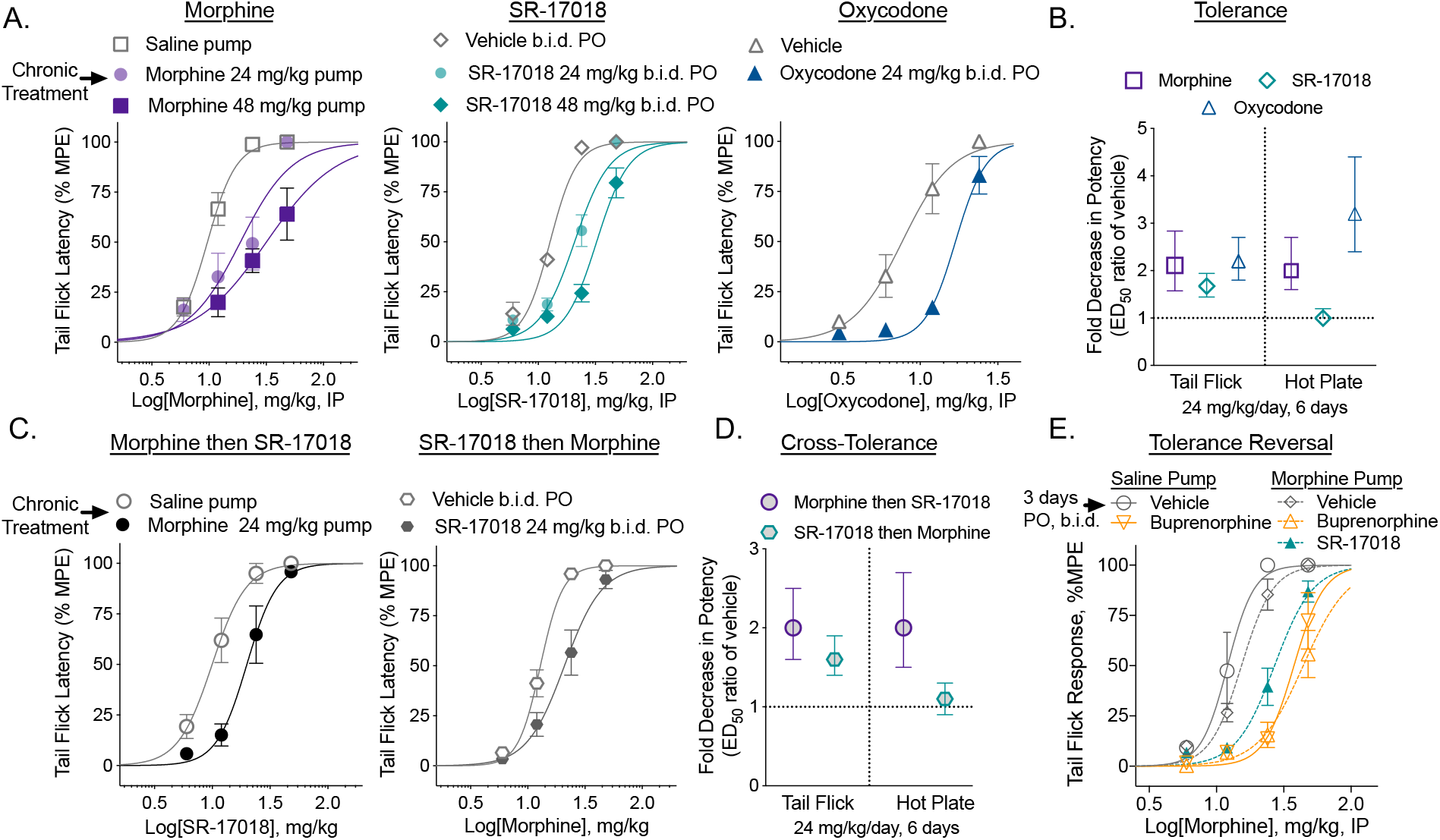
Chronic SR-17018 produces antinociceptive tolerance in the tail flick assay and morphine tolerance persists upon substitution with SR-17018 or buprenorphine in the tail flick assay. **A.** Male C57BL/6 mice were implanted with saline or morphine osmotic minipumps for 6 days or treated by PO, b.i.d. gavage with vehicle or SR-17018 or oxycodone. A cumulative (IP) dose response curve was performed on the 7^th^ day (12 hours after the last PO dose) (n= morphine: 12 saline; 7 at 24 & 48 mg/kg; SR17018: 8 vehicle; 10 at 24 & 48 mg/kg; oxycodone: 8 vehicle and 8 at 24 mg/kg; the mean with s.e.m. are shown). **B.** The shift in potency was compared between the saline treated and drug treated groups shown in A (presented as mean with 95% CI). Previously published data (Grim et al., 2019) for the hot plate potency shifts are included here for comparison. Confidence intervals that do no cross 1 are considered a significant shift in potency. **C.** Cross tolerance was determined as in A; wherein mice received 24 mg/kg/day of morphine (pump n=6) or 24 mg/kg/day SR-17018 (PO, b.i.d.) followed by a cumulative dosing paradigm to determine potency of the alternate compound (n = 6 all groups, mean, s.e.m. is shown). **D.** The shift in ED_50_ from C is presented as mean with 95% CI; the hot plate data are included from previously published data (Grim et al., 2019) as reference. **E.** Mice were treated with morphine (24 mg/kg/day) by osmotic pump for 6 days; then treated daily (PO, b.i.d.) with vehicle (n=13); SR-17018 (24 mg/kg/day, n=14) or buprenorphine (2 mg/kg/day, n=8) (by PO, b.i.d.) for 3 days, as indicated; saline pump treated (6 days) mice were treated daily (PO, b.i.d.) with vehicle (n=4) or buprenorphine (2 mg/kg/day, n=8). Tail flick response latency was then assessed following accumulative morphine dosing, 1 hour after each dose. Data are presented as mean ± s.e.m in A, C and E; with 95% CI in B and D; ED_50_ values with 95% CI are presented in Table 1.

**Table 1.** Potency of morphine, oxycodone and SR-17018 in the mouse tail flick assay (49°C warm water tail immersion) under acute and chronic treatment paradigms (means with 95% CI). ED_50_ are determined with maximum confined to 100% MPE.

| Treatment<br>Day 1 to 6 | Dose<br>mg/kg/day | Substitution<br>Day 7 to 10 | Dose<br>mg/kg/day | Challenge | ED<sub>50</sub> (95% CI) | ED<sub>50</sub> shift<br>(95% CI) |
| --- | --- | --- | --- | --- | --- | --- |
| Morphine | 0 | n/a | n/a | morphine | 9.7 (8.5 to 10.8) | n/a |
|  | 24 | n/a | n/a | morphine | 18.8 (13.6 to 25.3) | 2.1 (1.6 to 2.8) |
|  | 48 | n/a | n/a | morphine | 31.7 (22.7 to 57.8) | 3.4 (2.5 to 4.7) |
| SR-17018 | 0 | n/a | n/a | SR-17018 | 12.7 (11.6 to 13.7) | n/a |
|  | 24 | n/a | n/a | SR-17018 | 20.8 (18.4 to 23.4) | 1.7 (1.4 to 1.9) |
|  | 48 | n/a | n/a | SR-17018 | 32.4 (30.0 to 36.3) | 2.6 (2.2 to 3.0) |
| SR-17018<br>*females* | 0 | n/a | n/a | SR-17018 | 17.3 (15.3 to 19.4) | n/a |
|  | 48 | n/a | n/a | SR-17018 | 32.6 (26.5 to 39.5) | 1.9 (1.5 to 2.3) |
| Morphine | 0 | n/a | n/a | SR-17018 | 10.0 (8.4 to 11.7) | n/a |
|  | 24 | n/a | n/a | SR-17018 | 19.9 (16.6 to 23.7) | 2.0 (1.6 to 2.5) |
| SR-17018 | 0 | n/a | n/a | morphine | 12.9 (12.0 to 13.9) | n/a |
|  | 24 | n/a | n/a | morphine | 20.9 (17.6 to 24.8) | 1.6 (1.4 to 1.9) |
| Oxycodone | 0 | n/a | n/a | oxycodone | 7.7 (6.4 to 9.2) | n/a |
|  | 24 | n/a | n/a | oxycodone | 16.9 (15.3 to 18.6) | 2.2 (1.8 to 2.7) |
| Morphine | 0 | vehicle | 0 | morphine | 12.1 (9.8 to 14.6) | n/a |
|  | 0 | buprenorphine | 2 | morphine | 37.7 (32.2 to 43.7) | 3.1 (2.5 to 3.9) |
|  | 48 | vehicle | 0 | morphine | 15.4 (14.0 to 16.9) | 1.3 (1.1 to 1.6) |
|  | 48 | SR-17018 | 48 | morphine | 27.0 (24.2 to 30.2) | 2.3 (1.9 to 2.8) |
|  | 48 | buprenorphine | 2 | morphine | 44.1 (37.7 to 55.2) | 3.7 (2.9 to 4.6) |

For morphine tolerance reversal studies, all procedures and dosing were as described previously (Grim et al., 2019). Reported maximum possible effect (MPE%) from cumulative dosing is relative to tail flick baseline responses prior to starting cumulative dosing.

For repeated dosing in the paclitaxel-induced neuropathy model, hyperalgesia responses were measured on day 7 and drug dosing commenced on Day 8. Vehicle (b.i.d., PO); SR-17018 (48 mg/kg/day, b.i.d., PO) and oxycodone (24 mg/kg/day, b.i.d., PO) were administered every 12 hours from Day 8 through Day 11; von Frey thresholds were measured 1 hour after the morning dose (vehicle, 24 mg/kg SR-17018 or 12 mg/kg oxycodone) on Days 8 and 11. Criteria for exclusion were as described above; 1 mouse did not meet the baseline threshold; 1 mouse did not display >1g change in threshold on day 7.

### Analysis and statistics

Details regarding the number of animals, presentation of data and statistical analyses are presented in the individual figure legends and within the methods for each experimental paradigm. ED_50_ values with 95% confidence limits were generated by nonlinear regression analysis with the top constrained to 100% and the bottom constrained to 0% as the %MPE data plotted were normalized to baseline and the 30 seconds cutoff in the tail flick test using GraphPad Prism software (8.0). For the dose response curves that did not reach 100%, we have not performed a statistical comparison. To facilitate comparisons, potency ratios were generated to compare shifts in potency relative to chronic saline or vehicle conditions. A shift in potency was considered significant if the confidence limits did not include 1. For reversal of morphine tolerance studies, all comparisons were made to morphine response curves obtained in saline-pump-implanted mice, followed by pump explant and 3 days of twice daily vehicle dosing.

## Results

### Chronic administration of SR-17018 induces tolerance in the warm water tail immersion assay

Mice treated for 7 days with morphine (24 or 48 mg/kg/day, o.m.p.) developed tolerance as indicated by a 2-or 4-fold shift in morphine potency relative to mice treated with saline pumps for the same time (Figure 1A). Tolerance was also observed when SR-17018 (24 and 48 mg/kg/day PO, b.i.d.) or oxycodone (24 mg/kg/day PO, b.i.d.) was administered orally, 2 times per day (Figure 1A). The degree of tolerance (shift in the ED_50_ of subsequent corresponding opioid challenge) relative to the appropriate vehicle did not differ between the drugs when compared at 24 mg/kg daily which is in stark contrast to the observations made in the same mice assessed for hot plate responsiveness (hot plate responses were published (Grim et al., 2019); the average ED_50_ values with 95% CI are shown in Figure 1B for comparison). Female mice also developed tolerance to SR-17018 in the tail flick assay (SFig1); all potency comparisons can be found in Table 1.

When assessed for cross-tolerance, morphine-tolerant mice (7 days, 48mg/kg/day o.m.p. morphine) display tolerance to SR-17018 and SR-17018 tolerant mice display tolerance to morphine (Figure 1C). This contrasts the lack of cross tolerance observed between the two compounds in the hot plate assay using the same cohort of animals (Figure 1D). See Table 1 for all potencies and shifts in potency for tail flick data (mean with 95% CI). Tail flick and hot plate tests have previously shown dissimilar responses to morphine infusion, which might be attributed to differential effects of morphine on spinal and supraspinal sites. Assessing acute tolerance of morphine in both assays, leads to development of tolerance predominantly at the supraspinal level (hot plate test) but not at a spinal level (tail flick) (Langerman et al., 1995), which are opposite responses to our observations. Differential contributions of spinal and supraspinal sites may therefore be responsible for the development of tolerance across nociceptive assays. Ultimately, it is apparent that a single nociceptive testing paradigm in rodents may be inadequate to define a compound’s tolerance liability.

### Substitution with SR-17018 does not reverse morphine tolerance in the tail flick assay

Previously, we reported that treating morphine-tolerant mice for 3 days with SR-17018 restores morphine-potency when assessed in the hot plate assay (Grim et al., 2019). Here we show that morphine sensitivity is not restored in the tail flick assay upon subsequent treatment with SR-17018 or buprenorphine when given following chronic morphine (Figure 1E) (Grim et al., 2019). Buprenorphine has been shown to have antagonistic properties at high doses (Dum and Herz, 1981) and it is not clear if residual buprenorphine (12 hours from last dose of 1 mg/kg, PO) is causing the shift in the ED_50_ or if it is causing desensitization of the receptor; regardless, morphine is less efficacious following buprenorphine treatment of saline pump treated mice as well (Figure 1E), Table 1.

### Partial reversal of SR-17018 tolerance with PKC inhibitor

In order to gain mechanistic insight into the mode of action of antinociceptive tolerance we utilized chelerythrine, a potent protein kinase C inhibitor (Herbert et al., 1990). Previous studies implicate PKC-mediated regulation of MOR, particularly in spinal-reflex-mediated nociception (Bull et al., 2017; Granados-Soto et al., 2000; Mao et al., 1995; Narita et al., 1995; Smith et al., 1999). Since morphine tail flick tolerance in βarrestin2-KO mice could be reversed with chelerythrine (Bohn et al., 2002), we asked whether this inhibitor could reverse the tail flick antinociceptive tolerance observed following chronic SR-17018 treatment. In Figure 2, we show that repeated dosing of SR-17018 led to a decreased response to an acute challenge with SR-17018 (24 mg/kg) compared to vehicle (two-way RM-ANOVA, F_*(4, 64)*_ = 4.050, P=0.00550) and that this decrease could be partially reversed upon pretreatment with chelerythrine (two-way RM-ANOVA F_*(1, 15)*_ = 14.77, p=0.0016, (Figure 2A, B). However, chronic SR-17018 again retains efficacy in the hot plate test and this is not affected by pretreatment with chelerythrine (two-way RM-ANOVA P=0. 79) (Figure 2C, D).

**Figure 2.**
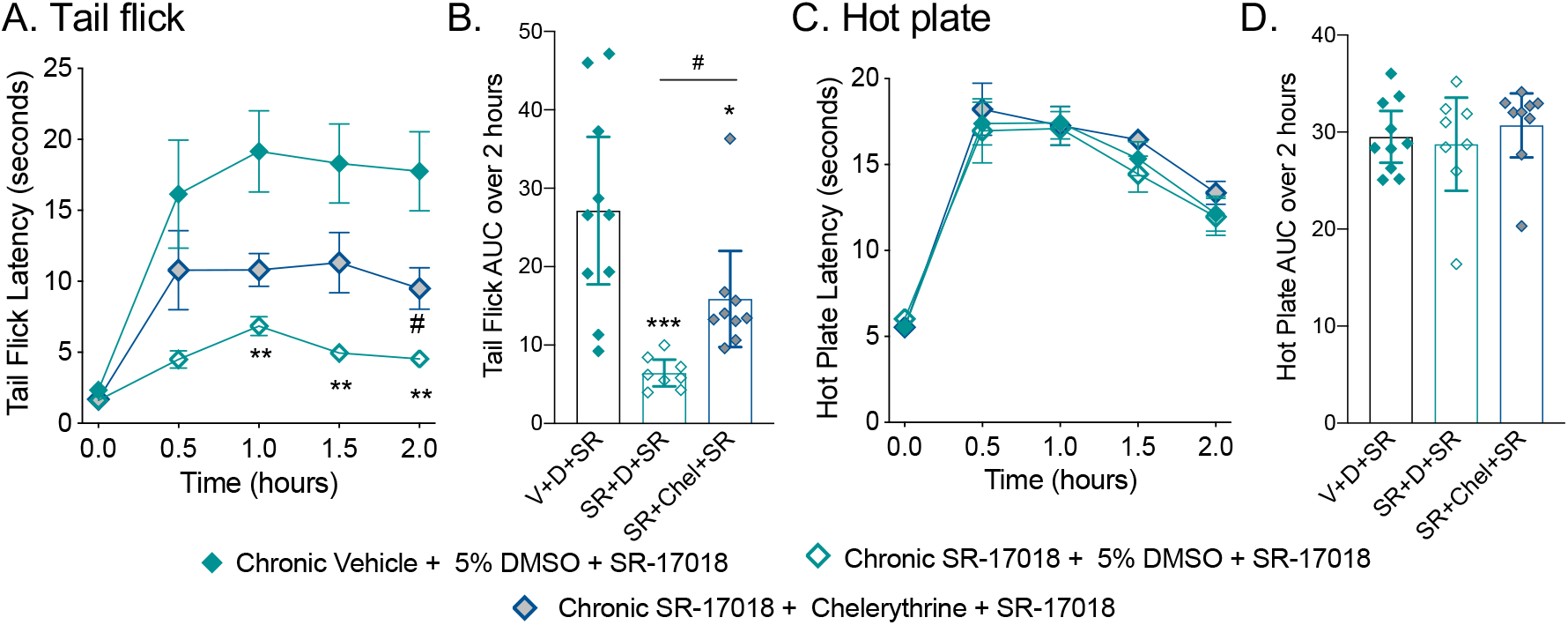
Chelerythrine partially reverses SR-17018-induced tolerance in the tail flick assay. A. Mice were treated as in Figure 1, with SR-17018 (24 mg/kg/day, PO, b.i.d.) or vehicle for 6 days. On day 7, mice were treated with 5% DMSO in water or chelerythrine (5 mg/kg, IP); 15 minutes later, they were tested for their response to SR-17018 (24 mg/kg, IP) in tail flick (A,B) and hot plate (C,D) tests at the times indicated. The area under the curve data are shown in B and D where V= chronic vehicle; D= 5% DSMO; SR= SR-17018; Chel = cheleryrthrine as indicated in the figure legends above. For A and C; data are presented as mean with s.e.m. Statistical comparisons in A are the result of a two-way ANOVA (interaction of time and treatment, p=0.0055) and Sidak’s post-hoc comparison (**p<0.01 for chronic vehicle + D + SR (n=10) vs. chronic SR + D + SR (n=8); ^#^p<0.05 for chronic SR + D + SR vs. chronic SR-+ Chel + SR (n=9). In B and D, data are presented as the mean with 95% CI; statistical comparison in B represents one-way ANOVA (p=0.0097); ***p<0.001 and *p<0.05 vs. V+D+SR; ^#^p<0.05 vs SR+D+SR.

### Comparison of opioid potency in the formalin assay

In order to further explore the utility of SR-17018 in another nociceptive pain model, a formalin response assay was employed to capture aspects of chemically induced acute pain (first phase, 0-10 min) and an inflammatory component of pain (second phase, 16-40 min) (Dubuisson and Dennis, 1977; Mogil et al., 1996; Tjolsen et al., 1992). Morphine, SR-17018 and oxycodone were evaluated for potency using single injections (IP) of each dose 45 minutes prior to formalin injection and compared to vehicle (Figure 3A, the same cohort of vehicle treated mice is graphed with each drug for comparison). The sum of the responses for each treatment were binned into the two phases and the ED_50_ values were calculated based on nonlinear regression following normalization of the data (Figure 3B). The three compounds do not differ in potency in either phase of the formalin assay based upon the co mparison of potencies with overlapping 95% CI.

**Figure 3.**
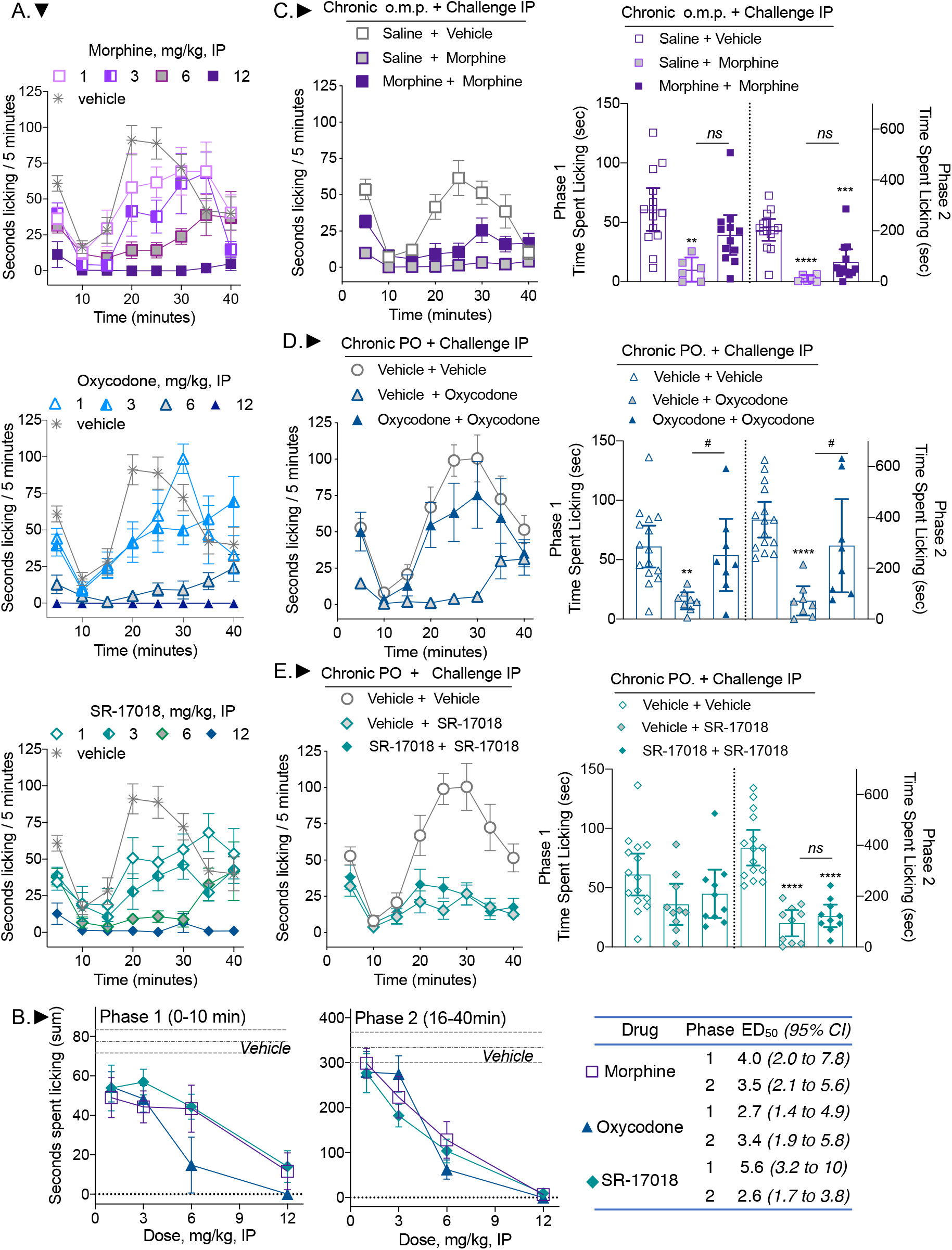
SR-17018 is potent in both phases of the formalin test and remains efficacious in the second phase of the formalin assay following chronic treatment. **A.** Mice were treated IP with vehicle (n=16), morphine (at 1, 3, 6 mg/kg, n=7 and at 12 mg/kg, n=5) oxycodone (at 1, 3 mg/kg, n=7 at 6 mg/kg, n=8 and at 12 mg/kg, n=3) or SR-17018 (at 1, 3, 6 mg/kg, n=7 and at 12 mg/kg, n=5). **B.** The sum of the time spent licking in each phase is reported per dose. The symbols are defined in the table. The mean ± s.e.m. are shown in the graphs. The ED_50_ calculated for each phase is presented with 95% CI. Mice were treated for 6 days with (**C**) saline pump followed by 9 mg/kg, IP morphine, n=6 or morphine 48 mg/kg/day followed by 9 mg/kg, IP morphine, n=12; (**D**) vehicle or oxycodone (24 mg/kg/day), PO, b.i.d. followed by 6 mg/kg IP oxycodone, n=8 for each group; (**E**) vehicle or SR-17018 (48 mg/kg/day) PO, b.i.d followed by 9 mg/kg IP SR-17018 challenge, n=10 per group. The vehicle + vehicle cohorts are the same in E and F. The mean sum of the time spent licking in both phases are presented in the left panels with 95% CI. Two-way RM-ANOVA was used to compare the effects over time (in the right panels) and the statistics are presented in the text. The comparison of sums was performed by one-way ANOVA where comparisons are: vs. saline + vehicle or vehicle + vehicle: **p<0.01; ***p<0.001; ****p<0.0001; vs. vehicle + oxycodone ^#^p<0.05.

### Evaluation of tolerance in the formalin assay

A separate cohort of animals was then treated for 6 days with morphine (48 mg/kg/day o.m.p), saline (o.m.p.), oxycodone (24 mg/kg/day, PO, b.i.d.), SR-17018 (48 mg/kg/day, PO, b.i.d.), or vehicle (PO, b.i.d.) as described for the thermal nociceptive studies in Figures 1–2. On the 7^th^ day, mice were administered an IP dose of the indicated compound 45 minutes prior to formalin injection. We found that the saline pump-implanted mice showed less of a response than mice treated acutely with vehicle (Figure 3A compared to 3C) and we could not determine if this was due to the surgical interference or if it was due to impaired access to reaching and licking their paws caused by the pumps implanted between the shoulders. Given that the response window was lower with saline pretreatment, some regard should be made towards the interpretations regarding the onset of morphine tolerance in this assay (Figure 3C); however, statistically, two-way RM-ANOVA indicates an interaction of time and treatment in phase 1 (F_*(2,29)*_=9.110, p=0.0009 and in phase 2 (F_*(8,116)*_=2.330, p=0.0234). A Tukey post-hoc analysis of phase one indicates that only the acute morphine group differs from the vehicle group in the first phase (p=0.0015); in the second phase, the acute morphine group differs from the vehicle control (p<0.0001) as well as the chronic morphine group (p=0.002) and the chronic morphine treatment group differs from vehicle control (p<0.0001). Since oxycodone is a similarly efficacious agonist as morphine and can be administered orally in the same manner as SR-17018, it serves here as a more relevant control than morphine. Repeated administration of oxycodone produces tolerance in both phases of the formalin assay (Figure 3D, two-way RM-ANOVA for time and treatment interaction: phase 1: F_*(2, 28)*_=4.057, p=0.0284, treatment effect phase 2: F_*(2, 28)*_= 12.53, p=0.0001). Acute oxycodone suppresses paw licking relative to vehicle in the chronic vehicle group (p=0.0036 phase 1, p<0.0001 phase 2) and relative to oxycodone treatment in the chronic oxycodone group (p=0.0345 phase 1, p=0.0158 phase 2). Following repeated administration of either vehicle or SR-17018, an acute dose of SR-17018 was not efficacious in the first phase of the test (Figure 3E, two-way RM-ANOVA, treatment effect: F_*(2, 32)*_=2.439, p=0.1033. However, acute SR-17018 effectively suppressed the response in the second phase following chronic vehicle (p<0.0001) or chronic SR-17018 (p<0.0001) demonstrating it retains efficacy following chronic treatment.

### Evaluation of opioids in a paclitaxel-induced neuropathy pain model

Finally, we asked whether SR-17018 has efficacy in another mouse model of pain, a chemotherapy-induced peripheral neuropathy model (Deng et al., 2012). Preclinical studies in rodents have shown that administration of paclitaxel results in long-lasting hypersensitivities to mechanical stimuli (Deng et al., 2015; Legakis et al., 2018; Smith et al., 2004). For the purpose of this study DBA/2J mice were used, as this strain exhibits especially robust changes in the neuropathic pain model (Smith et al., 2004). To induce hypersensitivity, mice were treated with paclitaxel as illustrated in the schematic (Figure 4A). Figure 4B shows the mean of individual responses with 95% CI of the baseline and days 7, 11 and 14 post-paclitaxel responses for all of the mice tested. All mice have developed robust hyperalgesia by day 7, which persists through day 14, as demonstrated by the significant decrease in the mechanical force threshold in comparison to baselines (Figure 4B; one-way RM-ANOVA, p<0.0001). The individual groups are shown with their drug treated cohorts; two-way RM-ANOVA reveals a significant interaction effect between treatment day and drug effect (vehicle or SR-17018 or morphine) (F_*(14, 146)*_ = 3.561, p<0.0001) followed by Tukey’s multiple comparison post hoc analysis (Figure 4C). SR-17018 increases the mechanical threshold tolerated at all of the doses tested mg/kg (Day 14 vs. 1 mg/kg, p=0.0067; 3 mg/kg, p=0.0010; 6 mg/kg, p=0.0019; 12 mg/kg, p=0.0039). Morphine has no effect at 1 mg/kg but was efficacious at 3 and 6 mg/kg (Day 14 vs. 1 mg/kg, p= 0.5784; 3 mg/kg, p=0.0196; 6 mg/kg, p=0.0055 (Figure 4C). Vehicle has no effect on treatment (day 14 vs. vehicle, p= 0.1297). The limited number of efficacious doses tested for morphine prevents an accurate determination of potency. For SR-17018, we determined the potency by hyperbolic curve fitting of the delta-delta (drug effect s – (day 14 s – baseline s), to be 2.1 (*0.45–6.9*) mg/kg, IP (ED_50_ with *95% CI*).

**Figure 4.**
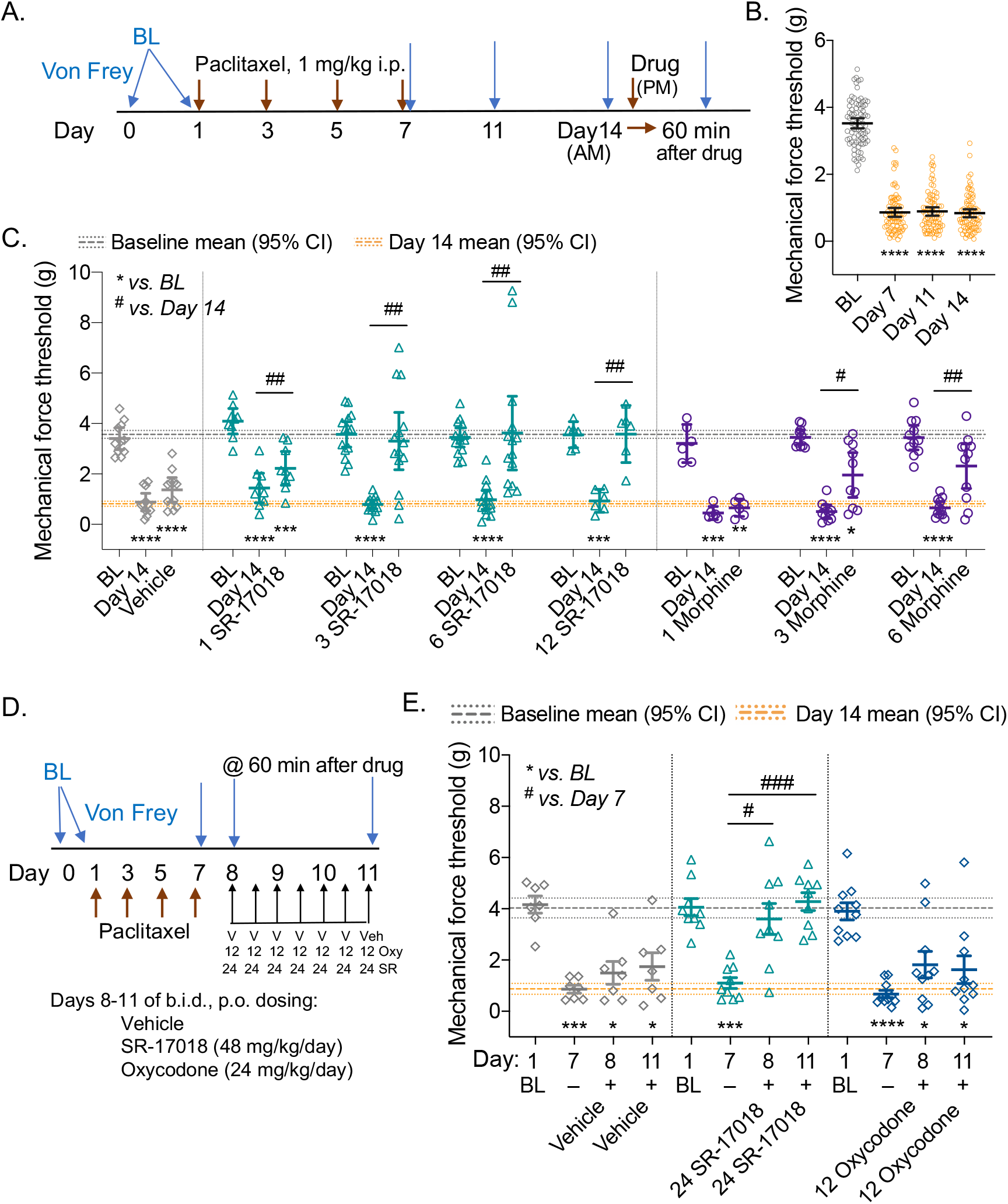
SR-17018 is effective in suppressing paclitaxel-induced neuropathic pain and retains efficacy upon repeated administration. **A.** Study design and timeline schematic. **B**. Mean of the baselines (BL) and Days 7, 11 and 14 thresholds recorded for all mice used in (C) with 95% CI (n=81). **C.** Baseline, Day 14 response and drug effect (test 1 h after drug) is presented for each paired cohort in the bottom panel wherein the dose is included as (mg/kg, IP) and the mean is presented with 95% CI; Student’s paired 2-tailed *t-*test: ****p<0.0001. The line on the ordinate at 3.6 g *(3.4–3.7)* is the mean BL and the line at 0.82 g *(0.71–0.92)* represents the mean of the Day 14 response *(95% CI)* for all cohorts from (B). Doses (mg/kg, IP) are indicated prior to the agonist, for the study. Vehicle: n=11, morphine: n=6 at 1, n=10 at 3, n=11 at 6 mg/kg; SR-17018: n=9 at 1, n=14 at 3, n=14 at 6, n=6 at 12 mg/kg. There is an interaction between measurement day and drug effect: two-way RM-ANOVA: F*_(14, 146)_* = 3.561, p<0.0001; Tukey’s multiple comparison post hoc analysis compared to BL ****p<0.0001, ***p<0.001, *p<0.05; Day 14 vs. drug effect: ^##^p<0.01; #p<0.05 within designated treatment groups. **D.** Study design timeline for chronic repeated dosing and testing. **E.** Baseline, Day 7 are compared to Day 8 and Day 11 where responses are measured 1 hour after dosing of Vehicle (PO, n= SR-17018 (24 mg/kg, PO) or oxycodone (12 mg/kg, PO). The line on the ordinate at 4.0 g *(3.6–4.4)* is the mean BL and the line at 0.87 g *(0.66–1.1)* represents the mean of the Day 7 response *(95% CI)* for all cohorts in this study. There is an interaction between measurement day and drug effect: two-way RM-ANOVA: F*_(6, 69)_* = 3.438, p=0.005; Tukey’s multiple comparison post hoc analysis compared to BL ****p<0.0001, ***p<0.001, *p<0.05; Day 7 vs. drug effect: ###p<0.001; ^#^p<0.05 within designated treatment groups.

### Evaluation of tolerance in paclitaxel-induced neuropathy pain model

To limit the duration of the test, we opted to begin chronic drug administration on Day 8 since hyperalgesia was evident by Day 7 (Figure 4B). Vehicle, SR-17018, and oxycodone were administered orally, every 12 hours (PO, b.i.d.) for 3 days and mechanical force thresholds were measured 1 hour after dosing on Days 8 and 11, as shown in the schematic (Figure 4D). There is an interaction of day and treatment effect (F_*(6, 69)*_ = 3.438, p=0.005, two-way RM-ANOVA); and vehicle did not elevate thresholds compared to Day 7 (Day 8: p=0.6386; Day 11: p=0.5614, Tukey post-hoc analysis) (Figure 4E). Compared to Day 7, SR-17018 (24 mg/kg, PO) was effective at elevating thresholds on Day 8 (p=0.0118) and after repeated dosing (Day 11: p=0.0004); oxycodone (12 mg/kg, PO) did not significantly alter threshold responses compared to Day 7 (Day 8: p=0.2344; Day 11: 0.3099).

## Discussion

In this study, we investigated a new opioid agonist, SR-17018, across several pain assays and assessed the development of tolerance upon repeated administration. SR-17018 produces tolerance in the warm water tail immersion test; this is in contrast to our previous results showing that these mice did not develop tolerance in the mouse hot plate test (Grim et al., 2019). We find that SR-17018 is efficacious a warm water tail immersion assay, a formalin paw-withdrawal assay and has similar potency as morphine. In the paclitaxel-induced neuropathy assay, we find that SR-17018 is more potent and efficacious than morphine and oxycodone. Furthermore, SR-17018 maintains efficacy in the second phase of formalin assay following a six day, twice daily dosing paradigm while oxycodone does not. In the paclitaxel-induced neuropathy pain model, SR-17018 demonstrates improved potency and efficacy over morphine and oxycodone; moreover, following a three-day twice daily dosing treatment paradigm, SR-17018 retains efficacy in this model as well.

We have previously shown that SR-17018 displays bias for G protein signaling over βarrestin2 recruitment in certain assays and it is attractive to speculate that this property underlies the favorable in vivo profile observed herein. However, we have not demonstrated that the decreased tolerance liability is directly due to the lack of βarrestin2 interactions in vivo. However, several studies have shown that disruption of βarrestin2 recruitment pathway dampens morphine tolerance (Bu et al., 2015; Li et al., 2009; Yang et al., 2011) including our own studies in the βarrestin2-KO mice (Bohn et al., 2002). Interestingly, the differential hot plate and tail flick responses to chronic SR-17018 are similar to outcomes observed in the βarrestin2-KO mice wherein they also developed morphine tolerance in the tail flick but not in the hot plate response. In an attempt to explore the mechanism underlying tail flick tolerance, we show that SR-17018 development of tolerance in the tail flick assay could be partially reversed by a PKC inhibitor; this approach also reversed morphine tolerance in the βarrestin2-KO mice (Bohn et al., 2002).

While βarrestin2 can regulate MOR in the spinal cord, many studies implicate other factors for regulating MOR in this system. Pharmacological inhibition of PKC, c-jun-N-terminal kinase and Src kinase have been shown to rescue morphine tolerance in the warm water tail immersion test in other studies (Bull et al., 2017; Granados-Soto et al., 2000; Marcus et al., 2015; Smith et al., 2002) and these mechanisms may ultimately contribute to the tolerance produced by agonists in the tail reflex nociception paradigm. Recent work in βarrestin2-KO mice that were backcrossed to C57BL6/J for the past 15+ years did not recapitulate the observations made in the mixed genetic background underscoring that MOR regulation is not entirely dependent on βarrestin2 (Kliewer et al., 2020; Kliewer et al., 2019). We reported in earlier studies that fentanyl and methadone produced hot plate tolerance in early generations of βarrestin2-KO mice (Raehal and Bohn, 2011) thereby demonstrating that the βarrestin-MOR interaction is not a universal mechanism underlying the development of opioid tolerance. Evaluation of MOR agonists with differential signaling properties, provide important tools for investigating nuances of MOR regulation while limiting the complications of mouse strain variations.

In addition to using the thermal nociception assays to characterize SR-17018 performance; additional pain models were also explored. Formalin injection into the paw pad captures two primary phases of response; the first phase is a display of a peripheral acute pain response driven by activation TRPA1 receptor and C-fibers, while the second phase captures an inflammatory component of pain accompanied by central sensitization of the dorsal horn (Hunskaar and Hole, 1987; Sanders et al., 2005; Savage and Ma, 2015; Tjolsen et al., 1992). The second phase is also influenced by supraspinal-mediated descending inhibition of spinal nociception (Basbaum and Fields, 1984; Detweiler et al., 1995; Manning and Franklin, 1998; Manning et al., 1994; Matthies and Franklin, 1992). Morphine has been shown to suppress responses in both phases of the test ((Martin et al., 2003; Sevostianova et al., 2003; Shannon and Lutz, 2002) and this study); and MOR-KO mice display an enhanced formalin response with a more robust second phase upon a 2% formalin injection (Zhao et al., 2003).

Demonstrating morphine antinociceptive tolerance in the formalin test has been more challenging (Abbott et al., 1981; Abbott et al., 1982; Connell et al., 1994; Detweiler et al., 1995). We were able to detect modest morphine tolerance in the second phase but not the first phase of the test. However, there were concerns that the implantation of the osmotic pump and the surgical interference made these results difficult to interpret, as the response to formalin following saline pump implantation was blunted overall. Oral delivery of oxycodone tolerance was evident in both phases of the assay following chronic treatment. Unfortunately, the 9 mg/kg dose of SR-17018 was not as efficacious in the first phase as anticipated from the acute dose response study following chronic vehicle treatment therefore, we cannot determine if tolerance developed in the first phase. The second phase of the formalin assay is considered to involve a supraspinal component as morphine tolerance in the second phase of the assay has been reported as the result of loss of supraspinal mediated descending inhibitory controls (Detweiler et al., 1995).

Using a model of chemotherapeutic-induced neuropathic pain we demonstrate that SR-17018 is more potent than morphine or oxycodone in suppressing allodynia that results from repeated paclitaxel injections. There are limited reports on the development of opioid tolerance in this chronic pain model and therefore we intended to look at tolerance following a six-day treatment period on Day 14. However, we noted that DBA mice showed weight loss during the chronic gavage regime (SR-17018: 85-93% and oxycodone: 93-96% of their initial weight) and displayed discomfort upon handling in the last days, independent of treatment group. Therefore, we did not compare responses on Day 14 opting to compare effects on Day 11 following the three day, twice-daily drug administration regimen. Oxycodone was used for comparison here since we hypothesized it would perform better than morphine and because we could orally administer it in the same vehicle as SR-17018. SR-17018 (24 mg/kg, PO) retained efficacy on the third day of chronic treatment (Day 11) compared to the first day of treatment (Day 8) although oxycodone (12 mg/kg, PO) did not significantly reverse allodynia on either day, although oxycodone is more potent than SR-17018 in the other pain assays (Figures 1, 3; Table 1). Since SR-17018 is more effective in alleviating paclitaxel-induced neuropathic pain than morphine and oxycodone; it is attractive to speculate that its interesting pharmacological properties contribute to this benefit.

SR-17018 was originally described as a biased agonist at the MOR (Schmid et al., 2017). Specifically, SR-17018 is nearly as potent as morphine in GTPγS assays in brainstem from mouse (SR17018: EC_50_= 288 ± 60; morphine: EC_50_=159±19 nM) and both are partial agonists ~40% Emax); yet SR-17018 produces very little βarrestin2 recruitment to the mouse MOR (SR17018: EC_50_> 10 μM; morphine EC_50_= 425 ± 51 nM) (Schmid et al., 2017). Upon comparing the performance of the two ligands, morphine and SR-17018 to DAMGO as a reference agonist within the same assays, SR-17018 has a bias factor of 102 and morphine is 1.9 (calculated from 10^ΔΔLogτ/K_A_). However, if the bias factors are calculated from the GTPγS binding in CHO cells expressing MOR, then the bias factor is 30 for SR-17018 and 0.80 for morphine. These comparisons demonstrate the contribution of context to the perception of bias when comparing compounds in different cellular models. For example, a recent study tested SR-17018 in a series of BRET-based signaling assays in transiently transfected cells and reported, based on their analysis that SR-17018 did not show bias (Gillis et al., 2020). This study also failed to reproduce biased signaling for other agonists that have been reported to show bias in other signaling platforms, including buprenorphine, oliceridine (TRV-130, Olinvyk^®^) and PZM21 (DeWire et al., 2013; Manglik et al., 2016; Ehrlich et al., 2019; Pedersen et al., 2020). In human trials, Olinvyk^®^ provides equal pain relief to morphine while presenting a significantly less postoperative drug-induced respiratory suppression (Ayad et al., 2020; Dahan et al., 2020). Ultimately, the effects of the drugs *in vivo* will determine the validity of the appropriate cellular signaling models for future drug development efforts.

As the development of MOR agonists is pursued in the attempt to refine the therapeutic potential and avoid unwanted side effects, it remains important to recognize that mechanisms by which a receptor signals and is regulated may not be conserved throughout all tissues and cells. Proximal location of regulation machinery may, in part, explain the differential responses that will depend on location of effect. In this study, we see a difference in adaptation upon chronic exposure that manifests as tolerance in the tail immersion assay; however, tolerance does not develop in other pain behavior assays. With these concepts in mind, it is important to thoroughly study new pharmacological probes and not make assumptions as to how they will act, simply based upon their characterization in binomial cellular signaling assays. We continue to seek understanding of how these ligands engage MOR to mediate their varied cellular responses. An agonist that can activate MOR in a manner to provide tolerance-resistant antinociception across several pain paradigms may ultimately be a beneficial therapeutic.

## Acknowledgements

Buprenorphine, and some morphine and oxycodone, were provided by the NIDA Drug Supply Program. We thank Ms. Nina McFague for additional assistance in blinded dosing and observations in the behavioral studies.

## Author contributions

FP and AAC-designing, performing and analyzing of the assays. Blinding, drug preparation, surgeries, dosing and scoring of the assays. Contributed to writing and editing of the manuscript.

TWG and CLS-designing, performing and analyzing behavioral studies; contributed to editing

NMK-synthesis of SR compounds

TDB-SR compound design and synthesis, funding and contributed to editing

LMB-experimental design, data analysis, funding and writing of the manuscript.

---

**Supplemental Figure 1.**
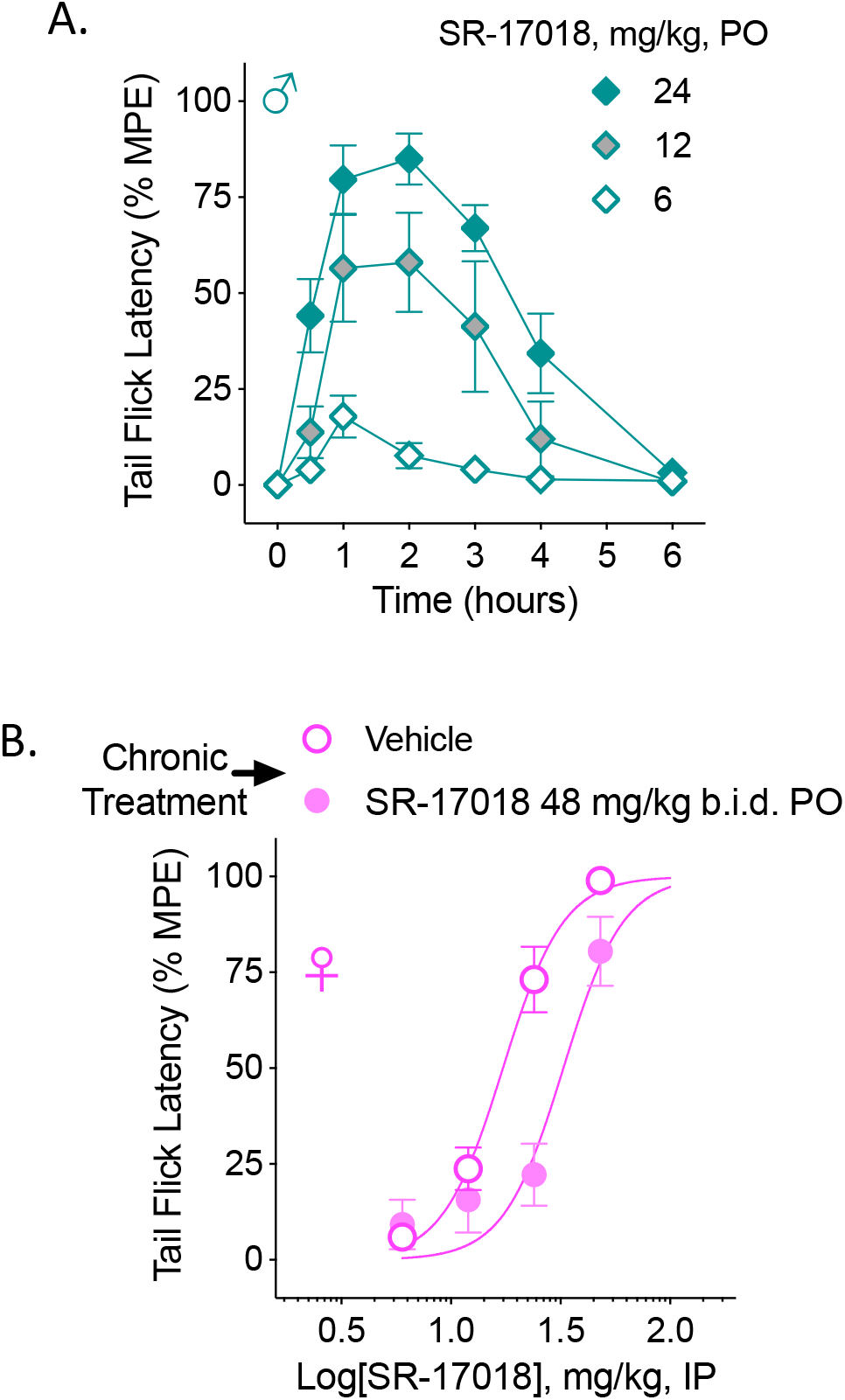
**A.** Male C57BL6/J mice: Dose response over time for oral (p.o.) dosing in the warm water (49°C) tail immersion assay (ED_50_: 11.1 *(6.4-19.0)* mg/kg, n=5 at 6, 12; n=8 at 24 mg/kg, PO). **B.** Female C57BL6/J mice: Dose response determined at 1 hour following intraperitoneal (IP) injection of SR-17018 following chronic treatment with vehicle (PO., b.i.d.) or 48 mg/kg/day SR-17018 (PO., b.i.d.) for 6 days. Potencies are provided in Table 1 of the text.

## References

Abbott, F. V., Franklin, K. B., Ludwick, R. J., Melzack, R., 1981. Apparent lack of tolerance in the formalin test suggests different mechanisms for morphine analgesia in different types of pain. Pharmacol Biochem Behav 15, 637–640.

Abbott, F. V., Melzack, R., Leber, B. F., 1982. Morphine analgesia and tolerance in the tail-flick and formalin tests: dose-response relationships. Pharmacol Biochem Behav 17, 1213–1219.

Ayad, S., Demitrack, M. A., Burt, D. A., Michalsky, C., Wase, L., Fossler, M. J., Khanna, A. K., 2020. Evaluating the Incidence of Opioid-Induced Respiratory Depression Associated with Oliceridine and Morphine as Measured by the Frequency and Average Cumulative Duration of Dosing Interruption in Patients Treated for Acute Postoperative Pain. Clin Drug Investig 40, 755–764.

Basbaum, A. I., Fields, H. L., 1984. Endogenous pain control systems: brainstem spinal pathways and endorphin circuitry. Annu Rev Neurosci 7, 309–338.

Bodnar, R. J., 2000. Supraspinal circuitry mediating opioid antinociception: antagonist and synergy studies in multiple sites. J Biomed Sci 7, 181–194.

Bohn, L. M., Gainetdinov, R. R., Lin, F. T., Lefkowitz, R. J., Caron, M. G., 2000. Mu-opioid receptor desensitization by beta-arrestin-2 determines morphine tolerance but not dependence. Nature 408, 720–723.

Bohn, L. M., Lefkowitz, R. J., Caron, M. G., 2002. Differential mechanisms of morphine antinociceptive tolerance revealed in (beta)arrestin-2 knock-out mice. J Neurosci 22, 10494–10500.

Bohn, L. M., Lefkowitz, R. J., Gainetdinov, R. R., Peppel, K., Caron, M. G., Lin, F. T., 1999. Enhanced morphine analgesia in mice lacking beta-arrestin 2. Science 286, 2495–2498.

Bu, H., Liu, X., Tian, X., Yang, H., Gao, F., 2015. Enhancement of morphine analgesia and prevention of morphine tolerance by downregulation of beta-arrestin 2 with antigene RNAs in mice. Int J Neurosci 125, 56–65.

Bull, F. A., Baptista-Hon, D. T., Sneddon, C., Wright, L., Walwyn, W., Hales, T. G., 2017. Src Kinase Inhibition Attenuates Morphine Tolerance without Affecting Reinforcement or Psychomotor Stimulation. Anesthesiology 127, 878–889.

Connell, B. J., Barnes, J. C., Blatt, T., Tasker, R. A., 1994. Rapid development of tolerance to morphine in the formalin test. Neuroreport 5, 817–820.

Dahan, A., van Dam, C. J., Niesters, M., van Velzen, M., Fossler, M. J., Demitrack, M. A., Olofsen, E., 2020. Benefit and Risk Evaluation of Biased mu-Receptor Agonist Oliceridine versus Morphine. Anesthesiology 133, 559–568.

Deng, L., Guindon, J., Cornett, B. L., Makriyannis, A., Mackie, K., Hohmann, A. G., 2015. Chronic cannabinoid receptor 2 activation reverses paclitaxel neuropathy without tolerance or cannabinoid receptor 1-dependent withdrawal. Biol Psychiatry 77, 475–487.

Deng, L., Guindon, J., Vemuri, V. K., Thakur, G. A., White, F. A., Makriyannis, A., Hohmann, A. G., 2012. The maintenance of cisplatin- and paclitaxel-induced mechanical and cold allodynia is suppressed by cannabinoid CB(2) receptor activation and independent of CXCR4 signaling in models of chemotherapy-induced peripheral neuropathy. Mol Pain 8, 71.

Detweiler, D. J., Rohde, D. S., Basbaum, A. I., 1995. The development of opioid tolerance in the formalin test in the rat. Pain 63, 251–254.

DeWire, S. M., Yamashita, D. S., Rominger, D. H., Liu, G., Cowan, C. L., Graczyk, T. M., Chen, X. T., Pitis, P. M., Gotchev, D., Yuan, C., Koblish, M., Lark, M. W., Violin, J. D., 2013. A G protein-biased ligand at the mu-opioid receptor is potently analgesic with reduced gastrointestinal and respiratory dysfunction compared with morphine. J Pharmacol Exp Ther 344, 708–717.

Dubuisson, D., Dennis, S. G., 1977. The formalin test: a quantitative study of the analgesic effects of morphine, meperidine, and brain stem stimulation in rats and cats. Pain 4, 161–174.

Due, M. R., Piekarz, A. D., Wilson, N., Feldman, P., Ripsch, M. S., Chavez, S., Yin, H., Khanna, R., White, F. A., 2012. Neuroexcitatory effects of morphine-3-glucuronide are dependent on Toll-like receptor 4 signaling. J Neuroinflammation 9, 200.

Dum, J. E., Herz, A., 1981. In vivo receptor binding of the opiate partial agonist, buprenorphine, correlated with its agonistic and antagonistic actions. Br J Pharmacol 74, 627–633.

Ehrlich, A. T., Semache, M., Gross, F., Da Fonte, D. F., Runtz, L., Colley, C., Mezni, A., Le Gouill, C., Lukasheva, V., Hogue, M., Darcq, E., Bouvier, M., Kieffer, B. L., 2019. Biased Signaling of the Mu Opioid Receptor Revealed in Native Neurons. iScience 14, 47–57.

Gillis, A., Gondin, A. B., Kliewer, A., Sanchez, J., Lim, H. D., Alamein, C., Manandhar, P., Santiago, M., Fritzwanker, S., Schmiedel, F., Katte, T. A., Reekie, T., Grimsey, N. L., Kassiou, M., Kellam, B., Krasel, C., Halls, M. L., Connor, M., Lane, J. R., Schulz, S., Christie, M. J., Canals, M., 2020. Low intrinsic efficacy for G protein activation can explain the improved side effect profiles of new opioid agonists. Sci Signal 13.

Granados-Soto, V., Kalcheva, I., Hua, X., Newton, A., Yaksh, T. L., 2000. Spinal PKC activity and expression: role in tolerance produced by continuous spinal morphine infusion. Pain 85, 395–404.

Grim, T. W., Schmid, C. L., Stahl, E. L., Pantouli, F., Ho, J. H., Acevedo-Canabal, A., Kennedy, N. M., Cameron, M. D., Bannister, T. D., Bohn, L. M., 2019. A G protein signaling-biased agonist at the mu-opioid receptor reverses morphine tolerance while preventing morphine withdrawal. Neuropsychopharmacology 45, 416–425.

Herbert, J. M., Augereau, J. M., Gleye, J., Maffrand, J. P., 1990. Chelerythrine is a potent and specific inhibitor of protein kinase C. Biochem Biophys Res Commun 172, 993–999.

Hunskaar, S., Hole, K., 1987. The formalin test in mice: dissociation between inflammatory and non-inflammatory pain. Pain 30, 103–114.

Johnson, J. L., Rolan, P. E., Johnson, M. E., Bobrovskaya, L., Williams, D. B., Johnson, K., Tuke, J., Hutchinson, M. R., 2014. Codeine-induced hyperalgesia and allodynia: investigating the role of glial activation. Transl Psychiatry 4, e482.

Kliewer, A., Gillis, A., Hill, R., Schmiedel, F., Bailey, C., Kelly, E., Henderson, G., Christie, M. J., Schulz, S., 2020. Morphine-induced respiratory depression is independent of beta-arrestin2 signalling. Br J Pharmacol 177, 2923–2931.

Kliewer, A., Schmiedel, F., Sianati, S., Bailey, A., Bateman, J. T., Levitt, E. S., Williams, J. T., Christie, M. J., Schulz, S., 2019. Phosphorylation-deficient G-protein-biased mu-opioid receptors improve analgesia and diminish tolerance but worsen opioid side effects. Nat Commun 10, 367.

Kline, R. H. t., Wiley, R. G., 2008. Spinal mu-opioid receptor-expressing dorsal horn neurons: role in nociception and morphine antinociception. J Neurosci 28, 904–913.

Langerman, L., Zakowski, M. I., Piskoun, B., Grant, G. J., 1995. Hot plate versus tail flick: evaluation of acute tolerance to continuous morphine infusion in the rat model. J Pharmacol Toxicol Methods 34, 23–27.

Legakis, L. P., Bigbee, J. W., Negus, S. S., 2018. Lack of paclitaxel effects on intracranial self-stimulation in male and female rats: comparison to mechanical sensitivity. Behav Pharmacol 29, 290–298.

Li, Y., Liu, X., Liu, C., Kang, J., Yang, J., Pei, G., Wu, C., 2009. Improvement of morphine-mediated analgesia by inhibition of beta-arrestin2 expression in mice periaqueductal gray matter. Int J Mol Sci 10, 954–963.

Manglik, A., Lin, H., Aryal, D. K., McCorvy, J. D., Dengler, D., Corder, G., Levit, A., Kling, R. C., Bernat, V., Hubner, H., Huang, X. P., Sassano, M. F., Giguere, P. M., Lober, S., Da, D., Scherrer, G., Kobilka, B. K., Gmeiner, P., Roth, B. L., Shoichet, B. K., 2016. Structure-based discovery of opioid analgesics with reduced side effects. Nature 537, 185–190.

Manning, B. H., Franklin, K. B., 1998. Morphine analgesia in the formalin test: reversal by microinjection of quaternary naloxone into the posterior hypothalamic area or periaqueductal gray. Behav Brain Res 92, 97–102.

Manning, B. H., Morgan, M. J., Franklin, K. B., 1994. Morphine analgesia in the formalin test: evidence for forebrain and midbrain sites of action. Neuroscience 63, 289–294.

Mao, J., Price, D. D., Phillips, L. L., Lu, J., Mayer, D. J., 1995. Increases in protein kinase C gamma immunoreactivity in the spinal cord of rats associated with tolerance to the analgesic effects of morphine. Brain Res 677, 257–267.

Marcus, D. J., Zee, M., Hughes, A., Yuill, M. B., Hohmann, A. G., Mackie, K., Guindon, J., Morgan, D. J., 2015. Tolerance to the antinociceptive effects of chronic morphine requires c-Jun N-terminal kinase. Mol Pain 11, 34.

Martin, M., Matifas, A., Maldonado, R., Kieffer, B. L., 2003. Acute antinociceptive responses in single and combinatorial opioid receptor knockout mice: distinct mu, delta and kappa tones. Eur J Neurosci 17, 701–708.

Martinov, T., Mack, M., Sykes, A., Chatterjea, D., 2013. Measuring changes in tactile sensitivity in the hind paw of mice using an electronic von Frey apparatus. J Vis Exp, e51212.

Matthies, B. K., Franklin, K. B., 1992. Formalin pain is expressed in decerebrate rats but not attenuated by morphine. Pain 51, 199–206.

Mogil, J. S., Kest, B., Sadowski, B., Belknap, J. K., 1996. Differential genetic mediation of sensitivity to morphine in genetic models of opiate antinociception: influence of nociceptive assay. J Pharmacol Exp Ther 276, 532–544.

Moriwaki, A., Wang, J. B., Svingos, A., van Bockstaele, E., Cheng, P., Pickel, V., Uhl, G. R., 1996. mu Opiate receptor immunoreactivity in rat central nervous system. Neurochem Res 21, 1315–1331.

Narita, M., Mizoguchi, H., Narita, M., Sora, I., Uhl, G. R., Tseng, L. F., 1999. Absence of G-protein activation by mu-opioid receptor agonists in the spinal cord of mu-opioid receptor knockout mice. Br J Pharmacol 126, 451–456.

Narita, M., Narita, M., Mizoguchi, H., Tseng, L. F., 1995. Inhibition of protein kinase C, but not of protein kinase A, blocks the development of acute antinociceptive tolerance to an intrathecally administered mu-opioid receptor agonist in the mouse. Eur J Pharmacol 280, R1–3.

Pedersen, M. F., Wrobel, T. M., Marcher-Rorsted, E., Pedersen, D. S., Moller, T. C., Gabriele, F., Pedersen, H., Matosiuk, D., Foster, S. R., Bouvier, M., Brauner-Osborne, H., 2020. Biased agonism of clinically approved mu-opioid receptor agonists and TRV130 is not controlled by binding and signaling kinetics. Neuropharmacology 166, 107718.

Raehal, K. M., Bohn, L. M., 2011. The role of beta-arrestin2 in the severity of antinociceptive tolerance and physical dependence induced by different opioid pain therapeutics. Neuropharmacology 60, 58–65.

Sanders, R. D., Giombini, M., Ma, D., Ohashi, Y., Hossain, M., Fujinaga, M., Maze, M., 2005. Dexmedetomidine exerts dose-dependent age-independent antinociception but age-dependent hypnosis in Fischer rats. Anesth Analg 100, 1295–1302, table of contents.

Savage, S., Ma, D., 2015. Experimental behaviour testing: pain. Br J Anaesth 114, 721–724.

Schmid, C. L., Kennedy, N. M., Ross, N. C., Lovell, K. M., Yue, Z., Morgenweck, J., Cameron, M. D., Bannister, T. D., Bohn, L. M., 2017. Bias Factor and Therapeutic Window Correlate to Predict Safer Opioid Analgesics. Cell 171, 1165–1175 e1113.

Sevostianova, N., Zvartau, E., Bespalov, A., Danysz, W., 2003. Effects of morphine on formalin-induced nociception in rats. Eur J Pharmacol 462, 109–113.

Shannon, H. E., Lutz, E. A., 2002. Comparison of the peripheral and central effects of the opioid agonists loperamide and morphine in the formalin test in rats. Neuropharmacology 42, 253–261.

Smith, F. L., Javed, R., Elzey, M. J., Welch, S. P., Selley, D., Sim-Selley, L., Dewey, W. L., 2002. Prolonged reversal of morphine tolerance with no reversal of dependence by protein kinase C inhibitors. Brain Res 958, 28–35.

Smith, F. L., Lohmann, A. B., Dewey, W. L., 1999. Involvement of phospholipid signal transduction pathways in morphine tolerance in mice. Br J Pharmacol 128, 220–226.

Smith, S. B., Crager, S. E., Mogil, J. S., 2004. Paclitaxel-induced neuropathic hypersensitivity in mice: responses in 10 inbred mouse strains. Life Sci 74, 2593–2604.

Tarselli, M. A., Raehal, K. M., Brasher, A. K., Streicher, J. M., Groer, C. E., Cameron, M. D., Bohn, L. M., Micalizio, G. C., 2011. Synthesis of conolidine, a potent non-opioid analgesic for tonic and persistent pain. Nat Chem 3, 449–453.

Tjolsen, A., Berge, O. G., Hunskaar, S., Rosland, J. H., Hole, K., 1992. The formalin test: an evaluation of the method. Pain 51, 5–17.

Yaksh, T. L., 1997. Pharmacology and mechanisms of opioid analgesic activity. Acta Anaesthesiol Scand 41, 94–111.

Yang, C. H., Huang, H. W., Chen, K. H., Chen, Y. S., Sheen-Chen, S. M., Lin, C. R., 2011. Antinociceptive potentiation and attenuation of tolerance by intrathecal beta-arrestin 2 small interfering RNA in rats. Br J Anaesth 107, 774–781.

Zhao, C. S., Tao, Y. X., Tall, J. M., Donovan, D. M., Meyer, R. A., Raja, S. N., 2003. Role of micro-opioid receptors in formalin-induced pain behavior in mice. Exp Neurol 184, 839–845.

